# Confidence-Enhanced Semi-supervised Learning for Mediastinal Neoplasm Segmentation ^1^

**DOI:** 10.1101/2024.07.22.604560

**Authors:** Xiaotong Fu, Shuying Zhang, Jing Zhou, Ying Ji

**Affiliations:** Center for Applied Statistics, Renmin University of China, Beijing, China; School of Statistics, Renmin University of China, Beijing, China; Department of Thoracic Surgery, Beijing Institute of Respiratory Medicine and Beijing Chao-Yang Hospital, Capital Medical University, Beijing, China

## Abstract

Automated segmentation of mediastinal neoplasms in preoperative computed tomography (CT) scans is critical for accurate diagnosis. Though convolutional neural networks (CNNs) have proven effective in medical imaging analysis, the segmentation of mediastinal neoplasms, which vary greatly in shape, size, and texture, presents a unique challenge due to the inherent local focus of convolution operations. To address this limitation, we propose a confidence-enhanced semi-supervised learning framework for mediastinal neoplasm segmentation. Specifically, we introduce a confidence-enhanced module that improves segmentation accuracy over indistinct tumor boundaries by assessing and excluding unreliable predictions simultaneously, which can greatly enhance the efficiency of exploiting unlabeled data. Additionally, we implement an iterative learning strategy designed to continuously refine the estimates of prediction reliability throughout the training process, ensuring more precise confidence assessments. Quantitative analysis on a real-world dataset demonstrates that our model significantly improves the performance by leveraging unlabeled data, surpassing existing semi-supervised segmentation benchmarks. Finally, to promote more efficient academic communication, the analysis code is publicly available at https://github.com/fxiaotong432/CEDS.

**Author summary:** In clinical practice, computed tomography (CT) scans can aid in the detection and evaluation of mediastinal tumors. The early detection of mediastinal tumors plays a crucial role in formulating appropriate treatment plans and improving patient survival rates. To reduce the high cost of manual annotation, researchers have attempted to employ convolutional neural networks (CNNs) for efficient automatic segmentation. However, the significant challenges arise due to the considerable variation in shape, size, and texture of mediastinal tumors, posing difficulties for the segmentation task. In this study, we introduce a confidence-enhanced module with a semi-supervised learning framework. By evaluating the model’s prediction confidence and selecting high-confidence predictions, we improve the efficiency and quality of data utilization. This approach demonstrates the achievement of accurate mediastinal tumor segmentation with only a minimal amount of labeled data. Our research not only provides an effective technical approach for automatic segmentation of mediastinal tumors but also opens up new possibilities for optimizing strategies in semi-supervised learning methods.

## Introduction

Mediastinal tumors are a common type of thoracic disease, characterized by tumors located in the central region of the chest, specifically in the mediastinum. The mediastinum is situated between the left and right pleural cavities and contains vital organs and structures such as the heart, esophagus, and trachea. Although the global incidence of mediastinal tumors is relatively low, it remains significant, with a prevalence rate ranging from approximately 0.77% to 1.68% [1–3]. Due to their specific location, even non-cancerous mediastinal tumors can cause severe health issues if left untreated, such as chest pain, severe myasthenia gravis, and aplastic anemia, significantly impacting human health and lifespan [4, 5]. Moreover, approximately 50% of patients with primary mediastinal tumors exhibit no significant clinical symptoms, which further complicates the timely detection, which are crucial in formulating appropriate treatment plans [6].

In recent years, with the rapid advancement of computed tomography (CT) technology, CT imaging has become a routine method for examining thoracic diseases. As an efficient, non-invasive imaging technique, CT imaging can provide clear images of the mediastinal region. Through CT scans, doctors can obtain information about the tumor’s location, size, and its surrounding tissues. In early research on CT image analysis, related tasks were primarily accomplished using radiomics-based methods combined with traditional machine learning techniques [7]. With the development and application of deep learning technology, more studies have begun to utilize convolutional neural network (CNN) models for medical image analysis. These models do not rely on domain-specific expertise or manually extracted features and have achieved significant results in medical image segmentation [8, 9]. Currently, the vast majority of related research has focused on developing fully supervised learning models based on large datasets for tasks such as breast nodules [10, 11], brain tumors [12, 13], lung nodules [14, 15] and organ segmentation [16–18]. However, research on mediastinal tumor segmentation is relatively scarce [19], especially in the context of limited annotated data.

Moreover, there are still some limitations in the mediastinal tumor segmentation research. First of all, mediastinal tumors present a wide range of locations, irregular shapes, and varying sizes, as illustrated in Fig. 1 (a) and (b). They often have indistinct boundaries and complex background information, making it difficult to establish accurate tumor segmentation models. Second, deep learning models typically require large amounts of well-annotated data for training, and obtaining such a large volume of high-quality annotated mediastinal tumor data is impractical due to the high cost. High-quality annotated data necessitates manual annotation at the pixel/voxel level by surgeons, requiring substantial expertise and time. For example, thoracic surgeons need to use specialized annotation software to delineate tumor regions layer by layer, with the annotation of each set of CT images taking an average of 30 to 40 minutes.

**Fig 1.**
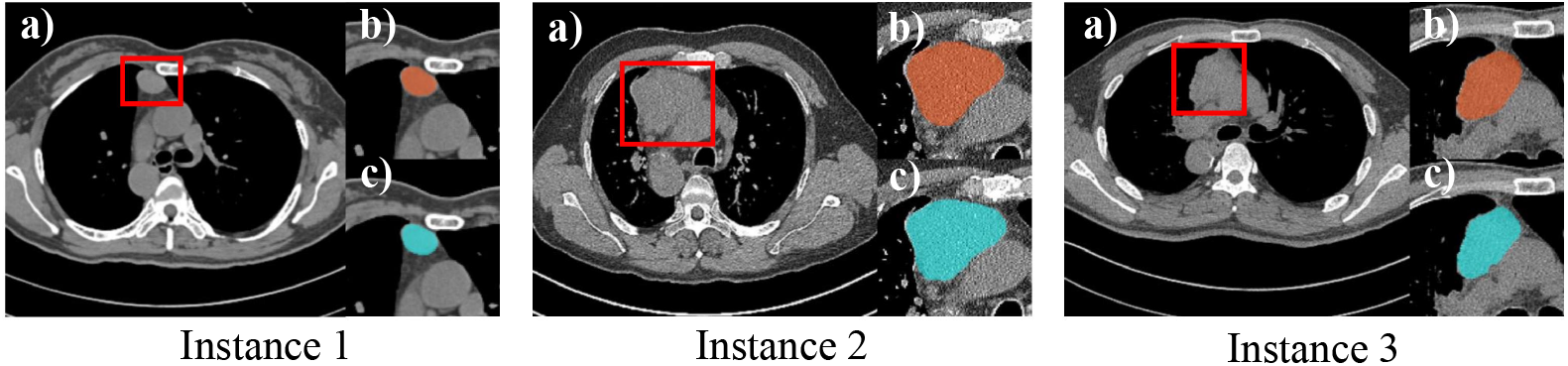
Three examples of mediastinal tumors with varying sizes and shapes. a) Original image with the tumor location highlighted; b) Zoomed-in view with tumor segmentation manually marked; c) Zoomed-in view with tumor segmentation generated by the model we have proposed in this article.

Given these constraints, we are inspired to develop an accurate semi-supervised segmentation model for mediastinal tumors, requiring only a minimal amount of labeled data. The semi-supervised learning (SSL) methods have become increasingly prevalent in medical image segmentation to mitigate the scarcity of manually annotated data. Researchers leverage the information of unlabeled data through various strategies, such as auxiliary tasks [20], data perturbation [21], and specially designed loss functions [22]. To the best of our knowledge, the exploration of using semi-supervised deep learning techniques for mediastinal tumor segmentation still remains untapped.

Motivated by the limitations of current studies, in this article, we propose an iterative confidence-enhanced semi-supervised learning framework specifically designed for segmenting mediastinal neoplasms. Specifically, we design an auxiliary decoder to estimate prediction confidence, which identifies more reliable pseudo labels to guide unsupervised learning. Considering the unique nature of the invasive boundaries of tumors, we further integrated an attention mechanism for better identification. To effectively optimize the proposed model, we also adopt an iterative training strategy to dynamically recalibrate the estimated confidence. Compared to existing research, our proposed method enhances the efficiency in utilizing unlabeled data, streamlines the training process, and is particularly effective in accurately delineating the subtle boundaries of mediastinal tumors. To summarize, our main contributions can be outlined as follows.

- First, we introduce a novel Confidence-Enhanced Dual-Decoder System (CEDS), a SSL model specifically designed for the precise segmentation of mediastinal tumors.
- Second, to improve the performance of the proposed model, we develop an iterative training strategy to ensure the system remain highly adaptive throughout the training process.
- Lastly, we demonstrate the efficacy of the proposed model through various experiments conducted on a real-world dataset, achieving a 1.44% improvement in dice coefficient compared to the benchmark methods, reaching a value of 88.62%.

## Related Work

In this section, we will first review the latest advancements and state-of-the-art methods in medical image segmentation. Next, we will focus on the progress of semi-supervised methods for lesion segmentation and discuss the shortcomings of existing research.

### Medical Image Segmentation

With the advancements of medical imaging technology, methods such as X-rays, CT scans, Magnetic Resonance Imaging (MRI), and ultrasound have become critical tools in clinical diagnosis and treatment [23]. Consequently, medical image segmentation is crucial for providing detailed visual information about diseases. This assists in the precise identification and extraction of key targets, such as organs or lesion areas, from images for further analysis and diagnosis. However, compared to traditional RGB images, medical images such as CT and MRI scans are typically grayscale, lacking the rich color information found in colored images. This makes it more difficult to distinguish different tissues and structures based on visual features [24]. Furthermore, the regions of interest in medical images are often of high similarity to surrounding tissues, where the boundaries are usually indistinct. Additionally, medical images are frequently affected by various artifacts (such as motion artifacts and noise), which further degrade image quality and increase segmentation difficulty. The variability in image acquisition conditions and equipment also introduces inconsistencies, requiring algorithms to have strong generalization capabilities to perform well across images from different devices [25]. These challenges make medical image segmentation still one of the most difficult tasks in the field of computer vision today.

Early approaches to medical image segmentation employed techniques such as edge detection [26], template matching [27], statistical shape models [28], active contour models [29], and early machine learning models [30]. These methods used artificially designed features to capture the boundary or structural information of images, achieving ideal results in certain scenarios. However, they often face challenges in effectively extracting useful features from medical images, especially when the images are blurred, noisy, or have low contrast.

Recently, the development of deep learning, particularly Convolutional Neural Networks (CNNs), has significantly accelerated progress in this field. These methods extract complex features automatically from large datasets without the need for manual feature design, significantly boosting both the accuracy and efficiency of segmentation. Such advancements have proven effective across various tasks, including breast nodule segmentation [31, 32], brain tumor segmentation [13, 33], lung nodule segmentation [15, 34], and organ segmentation [17, 18]. Initially, deep learning models mainly focused on slice-level 2D segmentation techniques aimed at pixel-level semantic classification. To achieve this, researchers developed end-to-end models based on encoder-decoder architectures, such as Fully Convolutional Networks (FCN) [35], U-Net [8], and Deeplab [36]. With the advancements in 3D imaging technology, models such as V-net [9], which build on the U-Net framework, have attracted attention for their superior performance in 3D segmentation tasks. These 3D segmentation algorithms better understand the spatial continuity of images, demonstrating enhanced performance when processing volumetric data. However, compared to 2D models, 3D algorithms require more computational resources due to an increase in parameters. To tackle this challenge, Kamnitsas et al. (2017) proposed DeepMedic [37], which uses a multi-scale 3D convolutional neural network to enhance segmentation accuracy while reducing computational complexity. Additionally, Isensee et al. (2021) introduced the nnU-Net [38], which retains the original U-Net structure but enhances it by incorporating adaptive preprocessing, model training, and inference strategies to create a more flexible learning framework. More recently, following the success of transformer models in natural language processing, Tragakis et al. (2023) introduced a fully convolutional transformer model [39], which combines the advantages of CNNs in image representation learning with the ability of transformers to capture long-term dependencies.

### Semi-supervised Segmentation

Semi-supervised learning (SSL) methods have become increasingly prevalent in medical image segmentation to address the scarcity of manually annotated data. Researchers leverage the information from unlabeled data through various strategies, such as auxiliary tasks [20], data perturbation [21], and specially designed loss functions [22].

The main semi-supervised methods can be roughly categorized as follows.

#### Pseudo labeling

The primary framework of early semi-supervised segmentation models was the pseudo-labeling method. This approach involves using pre-trained or fully supervised models to generate pseudo labels for unlabeled data. High-confidence pseudo labels are then added to the training dataset as annotated data, thereby increasing the data volume and improving model accuracy. This framework is highly flexible and can extend existing supervised models to semi-supervised scenarios. For instance, He et al. (2021) addressed pseudo label bias by employing distribution alignment and random sampling methods, ensuring they better match the true distribution [40]. Yuan et al. (2021) enhanced the efficacy of pseudo labeling by introducing a self-correcting loss that is resistant to noise [41]. Additionally, Yang et al. (2022) proposed the ST++ framework, which boosts model accuracy by selectively retraining based on the prediction reliability of unlabeled images [42].

#### Generative adversarial networks (GANs)

The generative adversarial networks (GANs) can simulate the distribution of real data in the training dataset and generate new data based on this distribution. A GAN model consists of two components: the generator and the discriminator. The generator creates images from random noise, mimicking the true distribution of the training dataset. The discriminator’s task is to distinguish between real images and fake images produced by the generator. This setup allows GANs to accurately replicate and learn from the data distribution of any dataset. The concept of GANs has been widely applied to tasks such as image generation [43], object detection [44], and semantic segmentation [45], and have been also extended to the field of semi-supervised segmentation. For example, Souly et al. (2017) used a generative network to provide fake training samples, optimizing the model by forcing real samples to be closer in feature space, thereby improving segmentation results [46]. Furthermore, Li et al. (2020) proposed SASSNET, which combined multi-task learning with the GAN concept, simultaneously predicting semantic segmentation and signed distance maps of object surfaces. In the meanwhile, they introduced adversarial loss during training to more effectively capture shape-aware features [47].

#### Consistency regularization

Finally, the consistency regularization methods are gaining popularity among researchers due to their simplicity, superior performance, and ability to be easily integrated with other frameworks. This method is based on the manifold or smoothness assumption, with the core idea that perturbations of data points within the same class should not alter the model’s output [48]. Researchers have explored various approaches to implementing consistency regularization. For instance, Tarvainen and Valpola (2017) introduced the mean-teacher framework, which calculates the consistency loss as the distance between the current model and a temporally averaged model [49]. Ouali et al. (2020) proposed cross-consistency training, which introduces model-level perturbations to enforce consistency between the predictions of the main decoder and those of the auxiliary decoders [50]. Meanwhile, Yang et al. (2023) presented the dual-stream perturbation technique UniMatch, where strongly perturbated views are simultaneously guided by a single weakly perturbated view [51]. However, such methods have certain limitations. If the model’s prediction for a data point is inaccurate, the consistency regularization process might treat this incorrect prediction as the “correct” learning target, introducing noise through consistency loss and negatively impacting the overall model performance. Therefore, more sophisticated strategies need to be designed to mitigate or avoid the negative effects of incorrect predictions on the model’s learning process. For example, Yu et al. (2019) quantified the uncertainty of model predictions by employing Monte Carlo sampling [52]. Kwon et al. (2022) have approached this by introducing an auxiliary error localization network to identify potentially erroneous pixels in pseudo-labels [53].

### Summary of Current Literature

Based on the above literature review, we found the current study faces several key challenges:

- Limited Existing Research: Few studies have explored the use of deep learning methods for mediastinal tumor segmentation. The rare existing studies typically rely on large volumes of densely annotated imaging data, which are costly to train and have limited generalization ability.
- High Manual Annotation Costs: Deep learning models generally require vast amounts of data for training. The high cost of acquiring large quantities of manually annotated data severely limits the practical clinical application of these models.
- Poor Segmentation Performance: Compared to organ segamentation, mediastinal tumors have indistinct boundaries, variable shapes, and complex background information, making it particularly challenging to develop accurate tumor segmentation models.

Therefore, the primary goal of this research is to develop a semi-supervised segmentation method for mediastinal tumors based on limited annotated data, which holds significant theoretical and clinical value.

## Methodology

### Overview

An overview of the proposed model is presented in Fig 2. Our goal is to improve the model’s representational capabilities through consistency learning with unlabeled data. Inspired by the mean teacher scheme, our approach employs both a teacher model and a student model [49]. The student model, which is used for training, progressively updates the teacher model using the exponential moving average (EMA) of its weights. This approach allows the teacher model’s predictions to serve as pseudo labels, guiding the learning process of the student model. To further enhance the model’s efficiency in utilizing unlabeled data, particularly in addressing indistinct tumor boundaries, we introduce the Confidence-Enhanced Dual-Decoder System (CEDS) into this framework. The CEDS features a shared encoder and dual decoders: one for segmentation and the other for confidence assessment. To customize the training process for the proposed architecture, we develop an iterative training strategy. This approach guarantees the dynamic adjustment of confidence estimation throughout the training process. Detailed explanations of each component are as follows.

**Fig 2.**
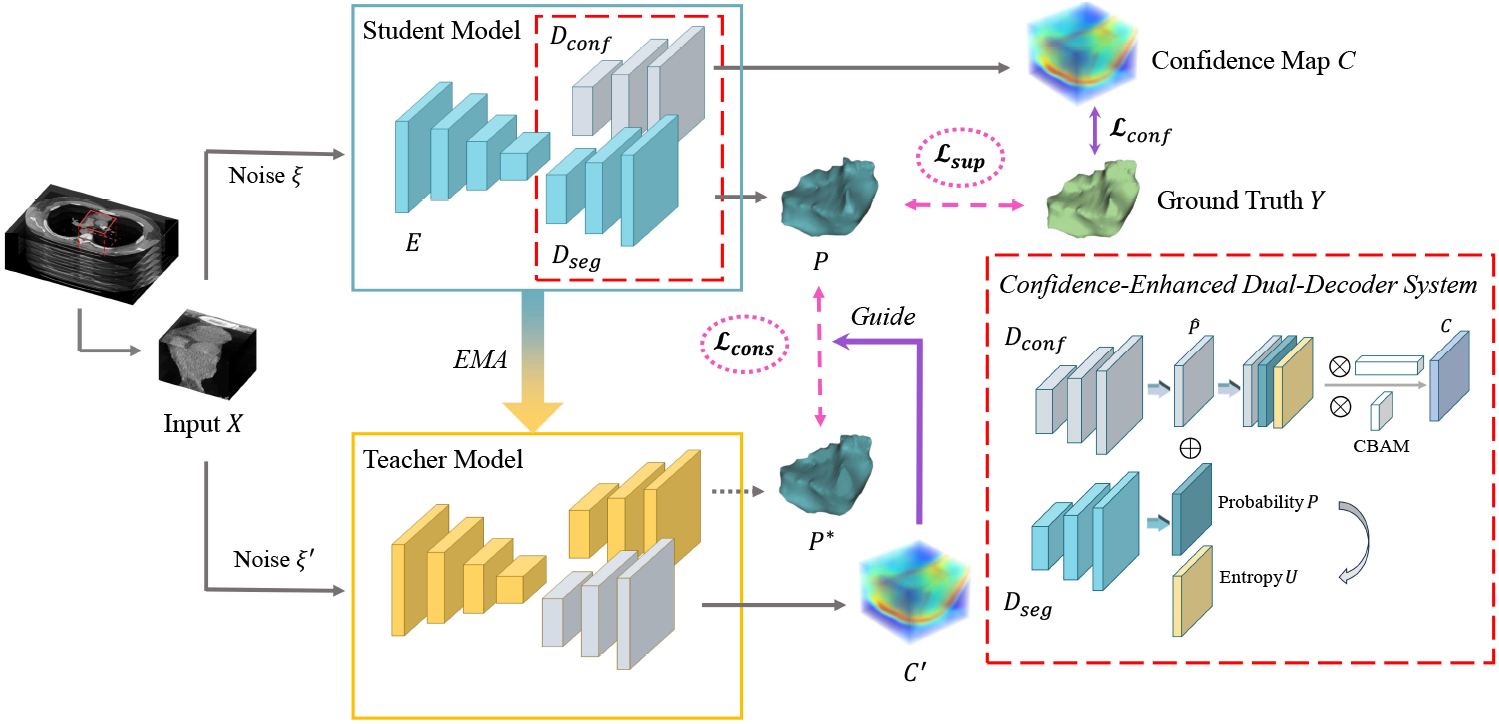
Overview of the proposed Confidence-Enhanced Semi-supervised Learning Framework for Mediastinal Neoplasm Segmentation. First, a 3D cropped block from a CT scan is fed into both the teacher and student models. Then, the teacher model, utilizing the EMA of the student model’s weights, generates segmentation predictions and a corresponding confidence map through two distinct decoders. Finally, this confidence map guides the calculation of consistency loss, which is determined by the differences in high-confidence predictions between the teacher and student models.

### Teacher-Student Semi-supervised Learning Scheme

In this study, we adopt the mean teacher framework, where the teacher model provides stable supervision to enhance the learning of the student model. Both of the two models are structured according to the idea of CEDS, which will be explained in the next section. Let *f* represent the student model and *f*′ be the teacher model, with their parameters denoted as *θ* and *θ*′, respectively. The student model updates *θ* using gradient descent, while the teacher model updates *θ*′ though the EMA of *θ*. Let *t* be the current step, then the updating process is defined as follows, where *α* is the hyperparameter regulating the update rate.

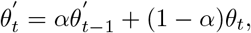

In this way, the weights of teacher network are averaged over multiple steps, which can reduce the noise and stabilize the training process. This improves the prediction of teacher model, providing more reliable targets for the student model and consequently enhancing overall performance.

Let *P* = *f* (*ξ*(*X*)) and *P*′ = *f*′ (*ξ*′ (*X*)) denote the segmentation predictions of the student model and teacher model for image *X*, respectively, with *ξ* and *ξ*′ representing the corresponding perturbations. Following the idea of Xie *et al*. [54], we transform *P*′ into soft pseudo labels *P*\* with a sharpening function (see Eq. 1), which is designed to integrate both consistency and entropy-minimization constraints.

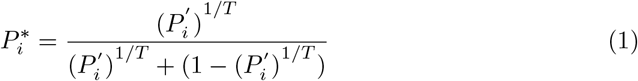

Here, *T* controls the sharpness; 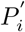 is the teacher model’s prediction at voxel *i*; and 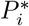 is its soft pseudo label. This transformation significantly enhances the efficiency of pseudo labels as a guidance.

The loss function of this scheme is composed of two main components. The first is the supervised loss ℒ_*sup*_ for the labeled dataset 𝒟_*L*_:

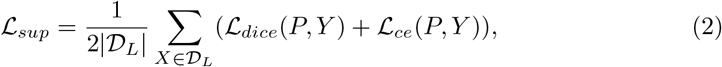

where ℒ_*dice*_ and ℒ_*ce*_ denote the standard dice and cross-entropy loss functions, respectively, with *Y* representing the ground truth labels. The second component is the unsupervised consistency loss ℒ_*cons*_, which is calculated as the mean squared error (MSE) between the predictions of the teacher model and student model across both 𝒟_*L*_ and the unlabeled dataset 𝒟_*U*_ (see Eq. 3). This approach is designed to maximize the utilization of available data.

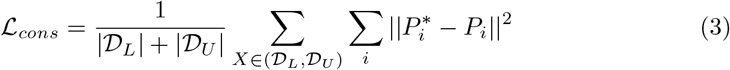

Then the total loss ℒ is computed as the sum of the supervised loss and the unsupervised consistency loss:

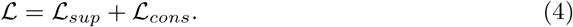

### Confidence-Enhanced Dual-decoder System

In this section, we will elaborate the structure of the proposed Confidence-Enhanced Dual-decoder System (CEDS). Employed by both the teacher and student models, the CEDS is designed to simultaneously generate segmentation predictions and assess their reliability, producing high-quality pseudo labels for subsequent training. To achieve this goal, existing methods typically involve an auxiliary network, such as the Error Correction Net (ECN), which identifies inaccurate predictions by comparing them with ground truths [55]. Although effective, they usually require a laborious two-stage training and fail to fully exploit the high-dimensional information of the segmentation network [53]. Therefore, we introduce the CEDS to simplify the training process and facilitate more accurate confidence estimations, especially for ambiguous segmentation boundaries.

#### CEDS Structure

The proposed CEDS has one shared encoder *E* and two specialized decoders: *D*_*seg*_ for segmentation and *D*_*conf*_ for confidence assessment, as highlighted in the red dash rectangle of Fig. 2. Specifically, *E* and *D*_*seg*_ replicate the V-net encoder-decoder architecture [9], while *D*_*conf*_ extends the V-net decoder with a Convolutional Block Attention Module (CBAM) to incorporate additional information. Here, the CBAM’s channel attention effectively integrates diverse features (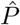, *P, U*), while its spatial attention mechanism enhances the extraction of spatial information from 3D CT images, thus improving the accuracy of confidence estimations for tumor boundaries. Let *P* = *D*_*seg*_(*ξ*(*X*)) denote the segmentation probability, and 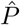 be the intermediate output from the V-net part in *D*_*conf*_. Note that 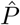 has the same shape as *P*. Subsequently, we calculate the voxel-level entropy map *U* from *P* to quantify prediction uncertainty. To integrate all of the information mentioned above, we employ the CBAM, denoted as *Atten*, to obtain the final confidence prediction

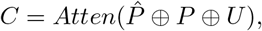

where ⊕ represents channel-wise concatenation. This architecture significantly reduces the model complexity without compromising its accuracy.

#### Loss Function

The segmentation network *D*_*seg*_ and *E* is trained using the supervised loss ℒ_*sup*_ on labelled data *D*_*L*_, which takes the same form as described in teacher-student scheme. Let *M* be the ground truth mask for confidence *C*, wherein each voxel *i* is assigned a value of 1 if the prediction *P*_*i*_ matches the ground truth *Y*_*i*_, and 0 otherwise:

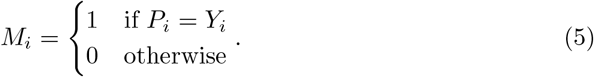

Then the loss for *D*_*conf*_, denoted as ℒ_*conf*_, is given by:

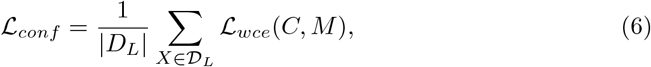

where ℒ_*wce*_ denotes the weighted cross-entropy loss function, with weights inversely proportional to class frequency. Note that the two decoders have two distinct training processes. The main segmentation network *D*_*seg*_ and *E* are updated together with ℒ_*sup*_.

In contrast, *D*_*conf*_ is independently updated with ℒ_*conf*_ to mitigate over-fitting and enhance efficiency. The initial few epochs employ ℒ_*sup*_ and ℒ_*conf*_ for model initialization.

Let 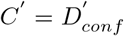 represent the confidence map generated by the teacher model. With *C*′, we can focus on higher-confidence predictions by selectively filtering out less reliable ones. The chosen predictions then serve as the learning targets for the student model. Thus, the confidence-enhanced consistency loss 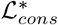 can be computed using the mean squared error (MSE) loss across valid voxels only:

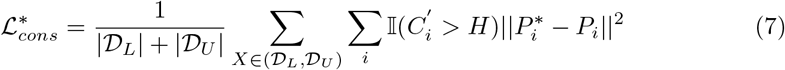

Here, 𝕀 is the indicator function, and *H* represents the threshold used to identify the most confident voxels. In the following experiments, we set it to 0.5 for simplicity. 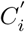 indicates the estimated confidence at voxel *i*. By replacing ℒ_*cons*_ with 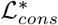 in Eq. 4, the confidence-enhanced total loss ℒ^*^ for CEDS can be formulated as

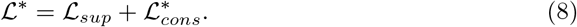

### Iterative Training Strategy

Note that we now have three parts of the proposed network: the shared encoder *E*, the segmentation decoder *D*_*seg*_ and the confidence decoder *D*_*conf*_. Considering the different convergence rates of the two decoders due to their distinct target tasks, especially with *D*_*conf*_ being prone to overfitting, we update the model with a special iterative training strategy. Specifically, it contains two phase: initialization and main training.

The initialization phase is critical for setting the model weights to appropriate initial values. During this phase, we use only the labeled data to train the network. For *D*_*seg*_ and *E*, we compute the supervised loss ℒ_*sup*_ and update the model accordingly. For *D*_*conf*_, we update its weights by calculating the loss ℒ_*conf*_, which is based on how accurate the predictions from the segmentation decoder *D*_*seg*_ are. Each decoder is updated once per epoch over a total of 𝒯_0_ epoch. Then in the main training phase, considering the different convergence rates of the two decoders due to their distinct target tasks, especially with *D*_*conf*_ being prone to overfitting, we update them at different rates. Specifically, we first use both labeled and unlabeled data to compute ℒ^*^ and update only *D*_*seg*_ and *E* for 𝒯_1_ epochs. Subsequently, we compute ℒ_*conf*_ and update *D*_*conf*_ independently for 𝒯_*conf*_ epochs. This process is then repeated. By employing this method, we can recalibrate *D*_*conf*_ with minimal additional training cost. Let *t* denote the current iteration and 𝒯_*conf*_ ≪ be the updating interval (with 𝒯_*conf*_ ≪ 𝒯_1_). Then the iterative training process is formulated as:

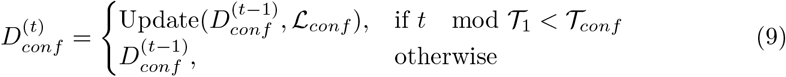

This training strategy helps to recalibrate *D*_*conf*_ based on the latest segmentation results, thereby improving the precision of confidence predictions. Moreover, since 𝒯_*conf*_ is set significantly smaller than 𝒯_1_ (i.e., less than 1%), this approach contributes only minimal additional training cost.

### Experiments

#### Dataset

We develop and evaluate the proposed model using a real-world dataset collected from a domestic hospital, and this study was approved by its ethics committee. The dataset consists of chest CT scans in NIfTI (nii.gz) format from 142 patients, which are randomly divided into a training set of 100 samples and a test set of 42 samples. Each sample includes a raw file of original images (197 to 556 slices, 512 × 512 pixels each) and a binary mask file with corresponding pixel-wise annotations. The mask is manually annotated by a surgeon with eight years of experience to ensure accuracy. The raw data undergoes a standard pre-processing pipeline [56, 57], which involves resampling to a uniform resolution of 0.625 × 0.625 × 0.625 mm per voxel, normalization of pixel values to the range [0,1], and foreground cropping centered around the lung region [58].

#### Implementation Details

We employ the stochastic gradient descent as the optimizer with a weight decay of 0.0001 and momentum of 0.9. The initial learning rate is set as 0.01, adjusted via cosine annealing schedule to a minimum of 0.001. A separate optimizer with the same setting is used for *D*_*conf*_. Training and updating intervals, 𝒯_1_ and 𝒯_*conf*_, are empirically determined to be 1000 and 5, respectively. The training process is set to 8,000 epochs to ensure convergence including an initialization phase of 400 epochs. The batch size is limited to two: one annotated and one unannotated image. For input, we randomly crop three sub-volumes of 112 × 112 × 80 from each image, maintaining a 2:1 ratio of positive (neoplasm-centered) to negative volumes to enhance training efficiency. To mitigate overfitting, we conduct data augmentation on the training set, including random flipping, rotation, and intensity shifting. All the experiments are conducted on a NVIDIA A30 GPU with 24GB of memory using PyTorch. For a comprehensive evaluation, we employ four metrics: dice coefficient, jaccard index, boundary accuracy (ACC), and boundary F1 score [19].

## Results

### Comparison Results with Benchmark Models

To validate the effectiveness of the proposed method, we conduct a comparative analysis against several well-established models: V-net [9], Uncertainty-aware Mean Teacher (UA-MT) [52], Shape-aware Strategy (SASSNet) [47], Dual-task Consistency (DTC) [59] and causality-inspired SSL (CauSSL) [60]. Note that V-net employs a fully supervised learning approach using exclusively labeled data, while the other models are semi-supervised learning methods.

Table 1 presents the quantitative results on our dataset. The proposed method achieves a 2.50% improvement in dice over the V-net trained on a limited labeled dataset (20% of the training set), approaching the performance of a fully supervised V-net using the complete labeled dataset (100% of the training set). Our model achieves the best performance among all semi-supervised models. Surprisingly, not all semi-supervised methods outperform the V-net given the same labeled dataset. This discrepancy can be attributed to the potential noise introduced by inaccurate pseudo labels, which consequently compromises model performance. Meanwhile, most SSL models are primarily developed for organ segmentation tasks, where the anatomical locations of organs in the images are relatively fixed. In contrast, mediastinal neoplasms, as a pathological condition, exhibit significant variability in their locations.

**Table 1.** Quantitative comparison results of supervised and semi-supervised methods. # Scans used: the number of CT scans treated as labeled or unlabeled in the training dataset, respectively.

| Method | # Scans used |  | Metrics |  |  |  |
| --- | --- | --- | --- | --- | --- | --- |
|  | Labeled | Unlabeled | Dice (%) | Jaccard (%) | ACC (%) | F1 score |
| V-Net (2016) | 20 | 0 | 86.12 | 81.32 | 90.08 | 0.7570 |
| V-Net | 100 | 0 | 90.85 | 86.13 | 92.52 | 0.8410 |
| UA-MT (2019) | 20 | 80 | 85.30 | 79.81 | 89.76 | 0.7630 |
| SASSNet (2020) | 20 | 80 | 85.23 | 79.86 | 90.28 | 0.7723 |
| DTC (2021) | 20 | 80 | 85.94 | 80.71 | 90.31 | 0.7685 |
| CauSSL (2023) | 20 | 80 | 87.18 | 82.21 | 90.33 | 0.7715 |
| Ours | 20 | 80 | **88.62** | **83.91** | **91.37** | **0.8005** |

Additionally, due to the invasive nature of tumors, they may infiltrate and adhere to surrounding organ tissues, resulting in more ambiguous boundaries. Therefore, directly transferring models designed for organ segmentation to this task results in a significant performance gap. In summary, the superiority of the proposed model primarily arises from two aspects. First, its specialized focus on mediastinal neoplasm segmentation, particularly emphasizing tumor segmentation from CT scans, a topic of limited attention in existing literature. Second, the incorporation of a confidence-enhanced segmentation strategy mitigates the effects of noise and maximizes the utility of information derived from unlabeled data.

Furthermore, to thoroughly evaluate the performance of each model in practical applications, we also conducted a detailed qualitative analysis, as presented in Fig. 1 c) and Fig. 3. Specifically, Fig. 1 c) shows the segmentation prediction generated by the proposed model on three examples, indicating the effectiveness of our model. Meanwhile, Fig. 3 displays performance comparisons of various SSL methods across three instances with distinct tumor morphologies. Overall, the proposed model exhibits superior performance in delineating tumor boundaries with high stability and accuracy, and effectively capturing the complex shapes of tumors for precise segmentation. However, constrained by the limited availability of labeled data, SSL methods still show some discrepancies between the predicted results and the true labels.

**Fig 3.**
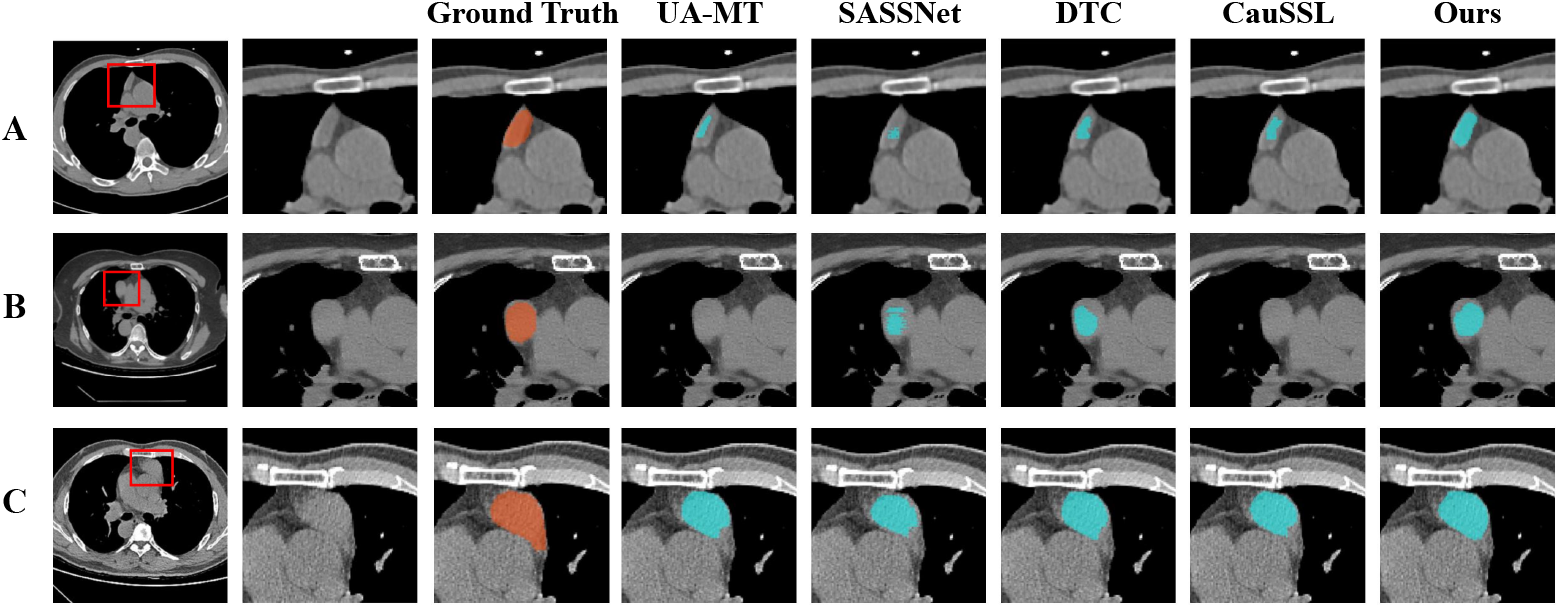
Qualitative comparisons of semi-supervised methods. Rows A to C display three different instances: first column, original CT images; second column, zoomed-in views highlighting the tumors; third column, manually annotated tumors; subsequent columns, tumor masks predicted by various semi-supervised methods.

### Ablation Study

To further evaluate the effectiveness of the key components in the proposed model, we conduct some ablation studies to validate the efficacy of each design of the proposed model. First, to assess the guidance effect of the confidence design, we bypassed the confidence estimation step and used all pseudo labels directly, reducing the model to a Mean Teacher model. Next, we evaluated the effectiveness of feature fusion by the attention mechanism in confidence estimation by removing the CBAM attention (*Atten*) from the CEDS structure. Finally, to determine whether the iterative training strategy better updates our proposed dual-decoder framework, we contrast it with a standard training approach. Table 2 shows the contributions of each component: confidence enhancement improves dice by 2.09%, CBAM by 1.26%, and iterative training by 0.35%. The outcomes validate the efficacy of our major designs.

**Table 2.**
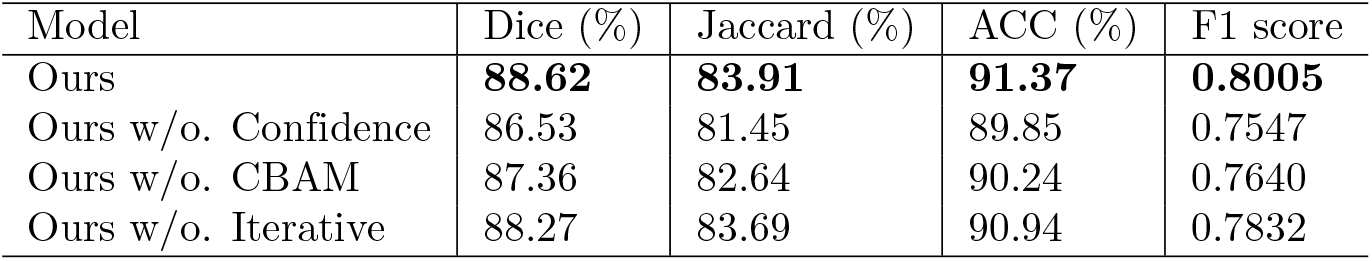
Ablation study on key designs of the proposed model.

### Visualization Analysis

To further explore the behavior of our model, we randomly selected an instance and visualized the corresponding confidence estimation as shown in Fig 4. Specifically, the main image on the left displays a heatmap of confidence around a specific tumor region. For better visualization, we present the uncertainty measure as 1 - confidence. Therefore, areas with higher temperatures indicate lower estimated confidence. This suggests exercising caution when using model predictions as pseudo-labels in these regions. Notably, voxels at the tumor boundaries exhibit lower confidence. This corresponds with general medical findings that due to the invasive boundaries of tumors, these areas are more prone to misclassification. This result also demonstrates our model’s capability to distinguish between reliable and unreliable segmentation results, enabling the selective incorporation of more accurate points into the calculation of consistency loss. This selective approach enhances the model’s overall accuracy and reliability in clinical settings. Additionally, we present three subplots on the right that illustrate the confidence distribution across different regions. From top to bottom, these subplots represent the distribution of tumor region, non-tumor regions, and the overall areas. The confidence levels significantly differ between the tumor area and non-tumor area, with generally lower confidence within the tumor area and higher confidence outside. Moreover, since the majority of the CT scan are non-tumor areas, the overall distribution closely resembles that of the non-tumor areas.

**Fig 4.**
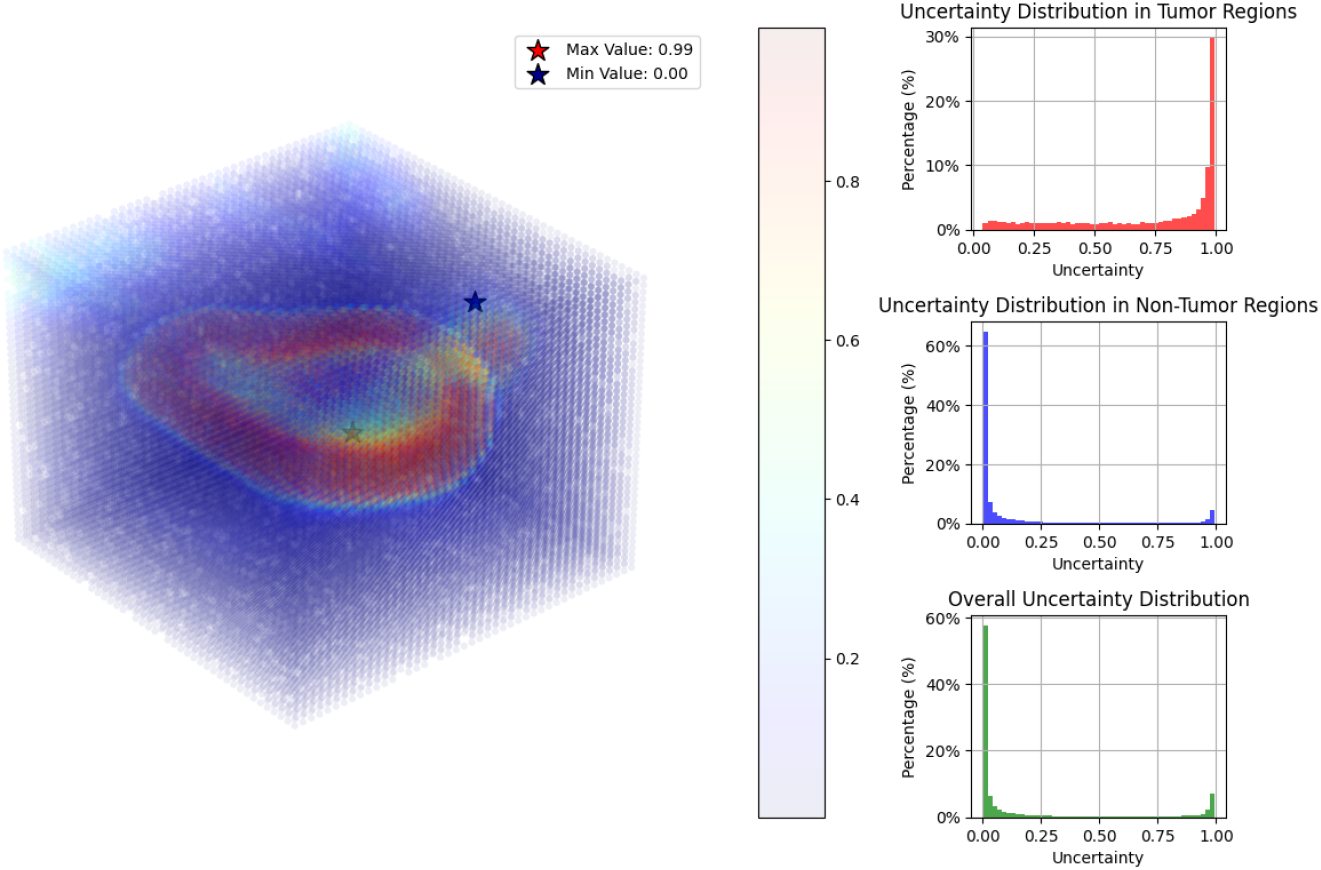
Visualization of estimated confidence map. The uncertainty is computed as 1-confidence. The main plot displays a heatmap centered on a tumor within a CT scan, with areas of higher uncertainty highlighted to show locations of lower model confidence. Star-marked points represent the maximum and minimum uncertainty values, at 0.99 and 0.00 respectively. The three subplots to the right quantify confidence distributions, specifically focusing on tumor regions (top), non-tumor regions (middle), and the overall scan (bottom), each illustrating the variability in model confidence across different anatomical contexts.

## Conclusion and Furture Work

### Conclusion

This study addresses the challenge of limited annotated data for mediastinal tumor segmentation by proposing a semi-supervised learning framework with confidence estimation. This framework not only fills the gap in current research, which often lacks focus on mediastinal tumor segmentation compared to organ segmentation, but also demonstrates the potential of leveraging unlabeled data to improve segmentation accuracy. By introducing the CEDS, we enhance the model’s performance in recognizing the blurry boundaries of tumors, achieving more precise tumor segmentation with the aid of unannotated data.

Specifically, our framework employs a teacher-student scheme to utilize unannotated data through temporal ensembling and consistency regularization. Additionally, we introduce a dual-decoder structure with an attention mechanism to assess the reliability of model predictions. This structure selectively uses more reliable predictions based on confidence, which helps reduce the negative impact of noise from unannotated data and effectively utilizes the information from such data. Additionally, we implement an iterative training strategy to optimally train the proposed structure. This strategy continuously recalibrates the estimated confidence in alignment with current segmentation predictions. Experimental results on a real-world dataset have validated the effectiveness of the proposed framework. Compared to traditional fully supervised learning methods and other semi-supervised learning strategies, our framework exhibits significant advantages in both quantitative evaluations and qualitative analysis.

The proposed model achieves a Dice coefficient of 88.62% and a boundary segmentation accuracy of 91.37%. Furthermore, additional ablation studies alos underscore the indispensable roles of confidence estimation, the attention mechanism, and iterative training in enhancing model performance.

### Future Work

Despite having achieved satisfied segmentation accuracy, this work still presents some certain limitations. Firstly, the model training and evaluation were limited to a specific mediastinal tumor dataset and have not been tested across other types of medical segmentation tasks. Therefore, the generalization ability of this framework to different medical segmentation tasks remains to be fully established. Secondly, some hyperparameters within the model, such as the reliability threshold *H* and the iterative update interval *T*_*conf*_, were chosen empirically based on experimental outcomes, without a systematic parameter search. There is potential for further enhancing model performance through a more detailed analysis of how these hyperparameters impact segmentation results. Lastly, the current model, implemented in Python, has not been converted into a tool or application that is easily operable by doctors or clinical practitioners. This limitation restricts its direct integration into clinical diagnostic processes, posing challenges to its practical application.

Considering these limitations, future work can expand in the following directions. First, to rigorously assess the model’s generalization ability, other widely used medical image segmentation datasets can be included for training and evaluation. Second, conducting a systematic parameter search may identify optimal hyperparameter settings. Finally, the model can be developed into user-friendly software, making it more accessible for clinical use.

## References

1. Henschke CI, Lee IJ, Wu N, Farooqi A, Khan A, Yankelevitz D, et al. CT screening for lung cancer: prevalence and incidence of mediastinal masses. Radiology. 2006;239(2):586–590.

2. Yoon SH, Choi SH, Kang CH, Goo JM. Incidental anterior mediastinal nodular lesions on chest CT in asymptomatic subjects. Journal of Thoracic Oncology. 2018;13(3):359–366.

3. Miyazawa R, Matsusako M, Nozaki T, Kobayashi D, Kojima F, Bando T, et al. Incidental mediastinal masses detected at low-dose CT screening: prevalence and radiological characteristics. Japanese Journal of Radiology. 2020;38:1150–1157.

4. Shahrzad M, Le TSM, Silva M, Bankier AA, Eisenberg RL. Anterior Mediastinal Masses. American Journal of Roentgenology. 2014;203(2):W128–W138.

5. Khatami A, Outhred AC, Britton PN, Huguon E, Lord DJ, Wong M, et al. Mediastinal mass in a healthy adolescent at The Children’s Hospital at Westmead, Australia. Thorax. 2015;70(2):194–197.

6. Davis Jr RD, Oldham Jr HN, Sabiston Jr DC. Primary cysts and neoplasms of the mediastinum: recent changes in clinical presentation, methods of diagnosis, management, and results. The Annals of thoracic surgery. 1987;44(3):229–237.

7. Zhao W, Xu Y, Yang Z, Sun Y, Li C, Jin L, et al. Development and validation of a radiomics nomogram for identifying invasiveness of pulmonary adenocarcinomas appearing as subcentimeter ground-glass opacity nodules. European journal of radiology. 2019;112:161–168.

8. Ronneberger O, Fischer P, Brox T. U-net: Convolutional networks for biomedical image segmentation. In: Medical Image Computing and Computer-Assisted Intervention–MICCAI 2015: 18th International Conference, Munich, Germany, October 5-9, 2015, Proceedings, Part III 18. Springer; 2015. p. 234–241.

9. Milletari F, Navab N, Ahmadi SA. V-net: Fully convolutional neural networks for volumetric medical image segmentation. In: 2016 fourth international conference on 3D vision (3DV). Ieee; 2016. p. 565–571.

10. Huang R, Lin M, Dou H, Lin Z, Ying Q, Jia X, et al. Boundary-rendering network for breast lesion segmentation in ultrasound images. Medical Image Analysis. 2022;80:102478.

11. Khaled R, Vidal J, Vilanova JC, Martí R. A U-Net Ensemble for breast lesion segmentation in DCE MRI. Computers in Biology and Medicine. 2022;140:105093.

12. Chen H, Qin Z, Ding Y, Tian L, Qin Z. Brain tumor segmentation with deep convolutional symmetric neural network. Neurocomputing. 2020;392:305–313.

13. Zhang D, Huang G, Zhang Q, Han J, Han J, Wang Y, et al. Exploring Task Structure for Brain Tumor Segmentation From Multi-Modality MR Images. IEEE Transactions on Image Processing. 2020;29:9032–9043.

14. Wang S, Zhou M, Liu Z, Liu Z, Gu D, Zang Y, et al. Central focused convolutional neural networks: Developing a data-driven model for lung nodule segmentation. Medical Image Analysis. 2017;40:172–183.

15. Singadkar G, Mahajan A, Thakur M, Talbar S. Deep deconvolutional residual network based automatic lung nodule segmentation. Journal of digital imaging. 2020;33:678–684.

16. Ngo TA, Lu Z, Carneiro G. Combining deep learning and level set for the automated segmentation of the left ventricle of the heart from cardiac cine magnetic resonance. Medical Image Analysis. 2017;35:159–171.

17. Gibson E, Giganti F, Hu Y, Bonmati E, Bandula S, Gurusamy K, et al. Automatic Multi-Organ Segmentation on Abdominal CT With Dense V-Networks. IEEE Transactions on Medical Imaging. 2018;37(8):1822–1834.

18. Huang H, Chen Q, Lin L, Cai M, Zhang Q, Iwamoto Y, et al. MTL-ABS3Net: Atlas-Based Semi-Supervised Organ Segmentation Network With Multi-Task Learning for Medical Images. IEEE Journal of Biomedical and Health Informatics. 2022;26(8):3988–3998.

19. Li J, Sun W, von Deneen KM, Fan X, An G, Cui G, et al. MG-Net: Multi-level global-aware network for thymoma segmentation. Computers in Biology and Medicine. 2023;155:106635.

20. Wang K, Zhan B, Zu C, Wu X, Zhou J, Zhou L, et al. Semi-supervised medical image segmentation via a tripled-uncertainty guided mean teacher model with contrastive learning. Medical Image Analysis. 2022;79:102447.

21. Bortsova G, Dubost F, Hogeweg L, Katramados I, De Bruijne M. Semi-supervised medical image segmentation via learning consistency under transformations. In: Medical Image Computing and Computer Assisted Intervention–MICCAI 2019: 22nd International Conference, Shenzhen, China, October 13–17, 2019, Proceedings, Part VI 22. Springer; 2019. p. 810–818.

22. Wu Y, Ge Z, Zhang D, Xu M, Zhang L, Xia Y, et al. Mutual consistency learning for semi-supervised medical image segmentation. Medical Image Analysis. 2022;81:102530.

23. Giger ML. Machine learning in medical imaging. Journal of the American College of Radiology. 2018;15(3):512–520.

24. Aljabri M, AlGhamdi M. A review on the use of deep learning for medical images segmentation. Neurocomputing. 2022;506:311–335.

25. Shen D, Wu G, Suk HI. Deep learning in medical image analysis. Annual review of biomedical engineering. 2017;19(1):221–248.

26. Yu-Qian Z, Wei-Hua G, Zhen-Cheng C, Jing-Tian T, Ling-Yun L. Medical images edge detection based on mathematical morphology. In: 2005 IEEE engineering in medicine and biology 27th annual conference. IEEE; 2006. p. 6492–6495.

27. Chen W, Smith R, Ji SY, Ward KR, Najarian K. Automated ventricular systems segmentation in brain CT images by combining low-level segmentation and high-level template matching. BMC medical informatics and decision making. 2009;9:1–14.

28. Tsai A, Yezzi A, Wells W, Tempany C, Tucker D, Fan A, et al. A shape-based approach to the segmentation of medical imagery using level sets. IEEE transactions on medical imaging. 2003;22(2):137–154.

29. Li C, Wang X, Eberl S, Fulham M, Yin Y, Chen J, et al. A likelihood and local constraint level set model for liver tumor segmentation from CT volumes. IEEE Transactions on Biomedical Engineering. 2013;60(10):2967–2977.

30. Li S, Fevens T, Krzyzżak A. A SVM-based framework for autonomous volumetric medical image segmentation using hierarchical and coupled level sets. In: International Congress Series. vol. 1268. Elsevier; 2004. p. 207–212.

31. Huang R, Lin M, Dou H, Lin Z, Ying Q, Jia X, et al. Boundary-rendering network for breast lesion segmentation in ultrasound images. Medical image analysis. 2022;80:102478.

32. Vidal J, Vilanova JC, Martí R, et al. A U-Net Ensemble for breast lesion segmentation in DCE MRI. Computers in Biology and Medicine. 2022;140:105093.

33. Chen H, Qin Z, Ding Y, Tian L, Qin Z. Brain tumor segmentation with deep convolutional symmetric neural network. Neurocomputing. 2020;392:305–313.

34. Wang S, Zhou M, Liu Z, Liu Z, Gu D, Zang Y, et al. Central focused convolutional neural networks: Developing a data-driven model for lung nodule segmentation. Medical image analysis. 2017;40:172–183.

35. Long J, Shelhamer E, Darrell T. Fully convolutional networks for semantic segmentation. In: Proceedings of the IEEE conference on computer vision and pattern recognition; 2015. p. 3431–3440.

36. Chen LC, Papandreou G, Kokkinos I, Murphy K, Yuille AL. Deeplab: Semantic image segmentation with deep convolutional nets, atrous convolution, and fully connected crfs. IEEE transactions on pattern analysis and machine intelligence. 2017;40(4):834–848.

37. Kamnitsas K, Ledig C, Newcombe VF, Simpson JP, Kane AD, Menon DK, et al. Efficient multi-scale 3D CNN with fully connected CRF for accurate brain lesion segmentation. Medical image analysis. 2017;36:61–78.

38. Isensee F, Jaeger PF, Kohl SA, Petersen J, Maier-Hein KH. nnU-Net: a self-configuring method for deep learning-based biomedical image segmentation. Nature methods. 2021;18(2):203–211.

39. Tragakis A, Kaul C, Murray-Smith R, Husmeier D. The fully convolutional transformer for medical image segmentation. In: Proceedings of the IEEE/CVF Winter Conference on Applications of Computer Vision; 2023. p. 3660–3669.

40. He R, Yang J, Qi X. Re-distributing biased pseudo labels for semi-supervised semantic segmentation: A baseline investigation. In: Proceedings of the IEEE/CVF International Conference on Computer Vision; 2021. p. 6930–6940.

41. Yuan J, Liu Y, Shen C, Wang Z, Li H. A simple baseline for semi-supervised semantic segmentation with strong data augmentation. In: Proceedings of the IEEE/CVF International Conference on Computer Vision; 2021. p. 8229–8238.

42. Yang L, Zhuo W, Qi L, Shi Y, Gao Y. St++: Make self-training work better for semi-supervised semantic segmentation. In: Proceedings of the IEEE/CVF Conference on Computer Vision and Pattern Recognition; 2022. p. 4268–4277.

43. Goodfellow I, Pouget-Abadie J, Mirza M, Xu B, Warde-Farley D, Ozair S, et al. Generative adversarial nets. Advances in neural information processing systems. 2014;27.

44. Bai Y, Zhang Y, Ding M, Ghanem B. Sod-mtgan: Small object detection via multi-task generative adversarial network. In: Proceedings of the European Conference on Computer Vision (ECCV); 2018. p. 206–221.

45. Luc P, Couprie C, Chintala S, Verbeek J. Semantic segmentation using adversarial networks. arXiv preprint arXiv:161108408. 2016;.

46. Souly N, Spampinato C, Shah M. Semi supervised semantic segmentation using generative adversarial network. In: Proceedings of the IEEE international conference on computer vision; 2017. p. 5688–5696.

47. Li S, Zhang C, He X. Shape-aware semi-supervised 3D semantic segmentation for medical images. In: Medical Image Computing and Computer Assisted Intervention–MICCAI 2020: 23rd International Conference, Lima, Peru, October 4–8, 2020, Proceedings, Part I 23. Springer; 2020. p. 552–561.

48. Chapelle O, Scholkopf B, Zien A Eds. Semi-Supervised Learning (Chapelle, O. et al., Eds.; 2006) [Book reviews]. IEEE Transactions on Neural Networks. 2009;20(3):542–542.

49. Tarvainen A, Valpola H. Mean teachers are better role models: Weight-averaged consistency targets improve semi-supervised deep learning results. Advances in neural information processing systems. 2017;30.

50. Ouali Y, Hudelot C, Tami M. Semi-supervised semantic segmentation with cross-consistency training. In: Proceedings of the IEEE/CVF conference on computer vision and pattern recognition; 2020. p. 12674–12684.

51. Yang L, Qi L, Feng L, Zhang W, Shi Y. Revisiting weak-to-strong consistency in semi-supervised semantic segmentation. In: Proceedings of the IEEE/CVF Conference on Computer Vision and Pattern Recognition; 2023. p. 7236–7246.

52. Yu L, Wang S, Li X, Fu CW, Heng PA. Uncertainty-aware self-ensembling model for semi-supervised 3D left atrium segmentation. In: Medical Image Computing and Computer Assisted Intervention–MICCAI 2019: 22nd International Conference, Shenzhen, China, October 13–17, 2019, Proceedings, Part II 22. Springer; 2019. p. 605–613.

53. Kwon D, Kwak S. Semi-supervised semantic segmentation with error localization network. In: Proceedings of the IEEE/CVF Conference on Computer Vision and Pattern Recognition; 2022. p. 9957–9967.

54. Xie Q, Dai Z, Hovy E, Luong T, Le Q. Unsupervised data augmentation for consistency training. Advances in neural information processing systems. 2020;33:6256–6268.

55. Mendel R, De Souza LA, Rauber D, Papa JP, Palm C. Semi-supervised segmentation based on error-correcting supervision. In: Computer Vision–ECCV 2020: 16th European Conference, Glasgow, UK, August 23–28, 2020, Proceedings, Part XXIX 16. Springer; 2020. p. 141–157.

56. Zhou J, Hu B, Feng W, Zhang Z, Fu X, Shao H, et al. An ensemble deep learning model for risk stratification of invasive lung adenocarcinoma using thin-slice CT. NPJ Digital Medicine. 2023;6(1):119.

57. Fu X, Meng X, Zhou J, Ji Y. High-risk Factor Prediction in Lung Cancer Using Thin CT Scans: An Attention-Enhanced Graph Convolutional Network Approach. In: 2023 IEEE International Conference on Bioinformatics and Biomedicine (BIBM); 2023. p. 1905–1910.

58. Hatamizadeh A, Tang Y, Nath V, Yang D, Myronenko A, Landman B, et al. Unetr: Transformers for 3d medical image segmentation. In: Proceedings of the IEEE/CVF winter conference on applications of computer vision; 2022. p. 574–584.

59. Luo X, Chen J, Song T, Wang G. Semi-supervised medical image segmentation through dual-task consistency. In: Proceedings of the AAAI conference on artificial intelligence. vol. 35; 2021. p. 8801–8809.

60. Miao J, Chen C, Liu F, Wei H, Heng PA. Caussl: Causality-inspired semi-supervised learning for medical image segmentation. In: Proceedings of the IEEE/CVF International Conference on Computer Vision; 2023. p. 21426–21437.

